# High-efficiency transfection of *Acanthamoeba castellanii* using a cationic polymer

**DOI:** 10.1101/2022.12.01.518696

**Authors:** Anaísa B. Moreno, Viktor Ek, Jens Eriksson, Mikael E. Sellin, Lionel Guy

**Affiliations:** Department of Medical Biochemistry and Microbiology, Science for Life Laboratory, Uppsala University, Uppsala, Sweden

**Keywords:** *Acanthamoeba*, transfection, cationic polymers, polyethylenimine, PEI

## Abstract

The free-living amoeba *Acanthamoeba castellanii* is an ecologically, clinically, and evolutionarily important microorganism. *A. castellanii* amoebae are directly pathogenic to humans and serve as reservoirs for bacterial pathogens (e.g., *Legionella pneumophila*) but also regulate the proliferation of other microorganisms in the soil. Despite their importance, no reliable genetic system has been developed, hampering the use of *A. castellanii* and related species as model organisms. Transfecting *A. castellanii* with plasmids is possible with commercial kits, but it is expensive, inefficient, and vulnerable to product discontinuation. In this contribution, we present a method for efficient transfection of *A. castellanii* with readily available and inexpensive polyethylenimines. We systematically explore the method’s parameters, obtaining up to 100-fold higher efficiency than currently used protocols. The method presented here provides a robust step towards a complete genetic toolbox for *A. castellanii*, hence expanding its use as a model organism.

## Introduction

*Acanthamoeba castellanii* is a free-living amoeba which is ubiquitous in various environments. It is a human pathogen responsible for two major infections – *Acanthamoeba* keratitis and granulomatous amoebic encephalitis (GAE) ^1–3^. It also serves as a reservoir for other microorganisms, such as bacteria, viruses and other protists, in either a parasitic or mutualistic relationship ^1^. *A. castellanii* has been used as a model organism to study cytoskeleton motility, host-pathogen interactions, gene expression, and the evolution of pathogenicity ^3,4^. However, this species is still underutilised relative to other amoebae, such as *Dictyostelium discoideum* ^3^. One of the reasons for this is its genomic complexity ^5,6^, which has hampered the development of molecular biology tools ^5,7^. The ability to effectively introduce and express exogenous genes in these amoebae is, thus, of particular interest and significance ^8^.

Transfection of *Acanthamoeba* with plasmid vectors has been successfully reported using physical methods (electroporation) ^9^, but more frequently with chemical methods, using the commercially available reagents SuperFect^®^ (Qiagen) ^10,11^, and Viafect™ (Promega) ^12^. However, these methods’ reported transfection efficiency is low, typically 5% or less ^2,7^. Commercial kits offer a quick and straightforward technique for transfection but are costly, which restricts the freedom to optimise the method for specific applications. Moreover, kits may be suddenly discontinued, as the example of SuperFect^®^ shows. Since the components for such reagents are patented, finding a replacement is difficult.

Cationic polymers such as polyethylenimines (PEIs), poly-L-lysines (PLLs), polyamidoamines (PAMAMs), and chitosans have been employed as gene delivery systems in eukaryotic cells ^13–15^. With their high number of positively charged (e.g., amines) residues, they form polyplexes with the negatively charged phosphate groups in DNA molecules via electrostatic interactions ^15^. This complexation condenses the DNA into small, compact particles, which cells internalise, and make their way to the nucleus where genes can be expressed ^14,15^. PEIs are among the best-studied cationic polymers due to their high cation charge density, wide availability, and low-cost ^14,16,17^. In addition, their amine groups can be polymerised in two ways to create linear or branched isomers. The degree of polymerisation, and branching of PEIs, along with other physio-chemical properties, affect their molecular weight, polydispersity, and buffering capacity, reflected in how well they interact with DNA. Although PEIs have been established as DNA carrier vectors for a variety of cell types ^15,17^, transfection efficiency may be affected by the nucleic acid quantity and purity, the preparation and composition of the complexes and the transfection conditions employed ^14,17,18^.

While PEIs have been used to transfect another free-living amoeba – *Naegleria* ^19^, they remain unexplored for *Acanthamoebae*. In this contribution, we describe the use of PEIs for high-efficiency transfection of *A. castellanii*. We systematically explore three types of PEI and assess their effect on cell toxicity and transfection efficiency. Importantly, we show that the here presented PEI-based protocol is vastly superior to current commercial standards for the transfection of *A. castellanii*. This provides a foundation for developing a full genetic system for this ecologically important amoeba.

## Results

### DNA condensation by PEI is affected by the complexation media

We first investigated how different complexation media and PEI amounts influenced the condensation of DNA. The media tested were selected for supporting *A. castellanii* growth – PYG, Ac medium and LoFlo, or for having previously been described in the literature as buffers used for PEI transfections – HBS, NaCl, and ultrapure water. Agarose-gel retardation assays were performed to evaluate the DNA condensation ability of the PEIs. After gel electrophoresis, the migration of the polyplexes was compared to that of naked plasmid DNA (pDNA) (**Figure 1A and S1**). In all conditions, increasing concentrations of PEI led to better complexation of DNA. However, the media used affected the complexation, most noticeably comparing the linear PEIs (PEI-25 and PEI-40) to the branched PEI (PEI-Br) (**Figure S1**). Nevertheless, a high stoichiometry of PEI to pDNA resulted in successful complexation across all buffer conditions. Retardation of the polyplexes can be due to the neutralisation of the negatively charged DNA (which would then not move through the gel), polyplex size change, or a combination of both. Overall, PYG appeared to work best with all PEIs (**Figure 1A**, compared to **Figure S1**), and since this is also the preferred medium for *A. castellanii*, PYG was selected for the remaining studies.

**Figure 1.**
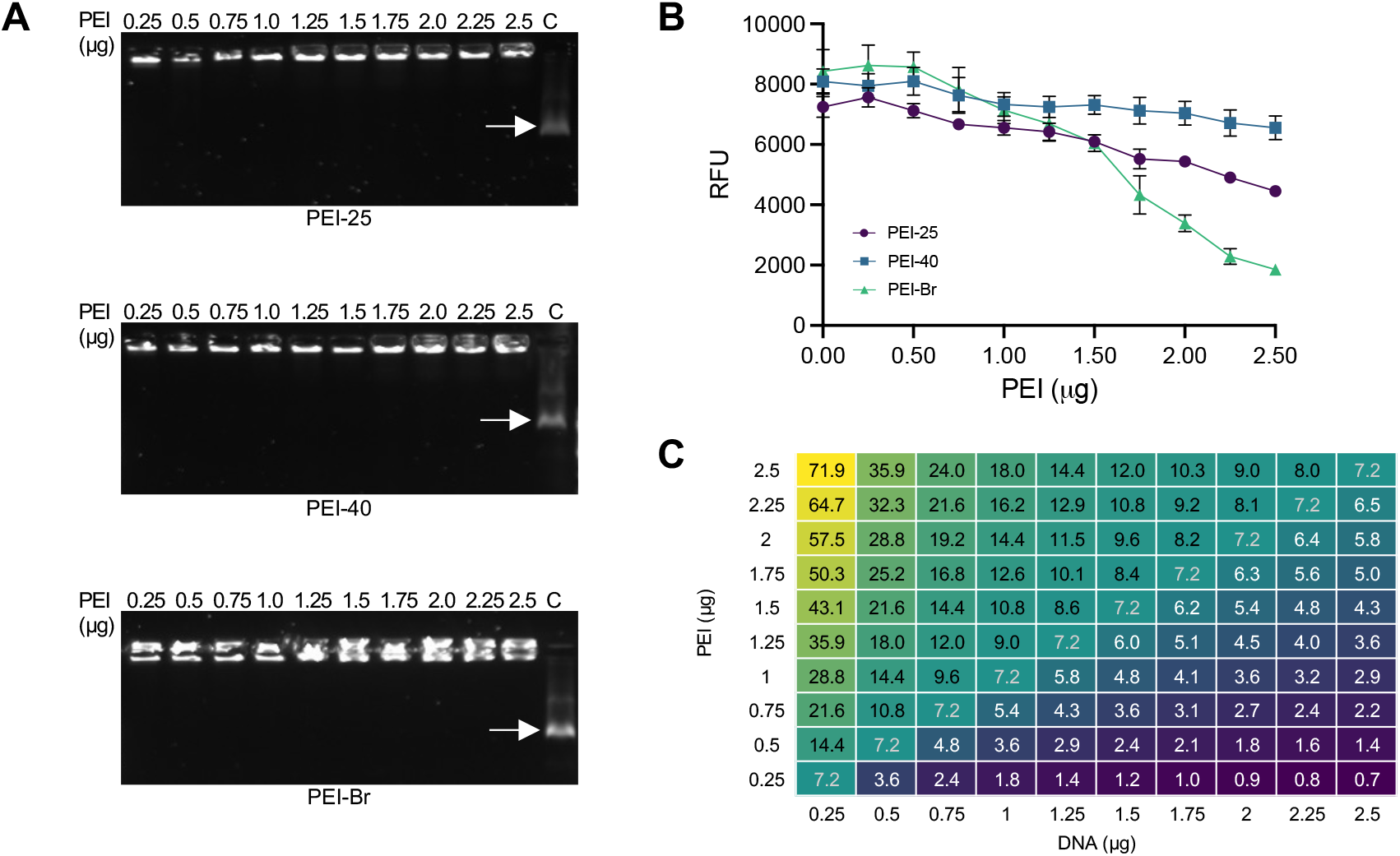
Condensation of PEI in PYG, cytotoxicity, and optimisation of transfection conditions. **(A)** DNA condensation ability of PEI in PYG. Increasing amounts of PEI-25, PEI-40 and PEI-Br were used to form complexes with 0.25 μg of pDNA. The migration of the pDNA-PEI polyplexes was compared to that of non-complexed pDNA, indicated by the white arrows (C, untreated control). **(B)** Cytotoxicity of PEI on *A. castellanii*. Viability of *A. castellanii* with increasing amounts of PEI in the medium was evaluated by resazurin assay. Resorufin fluorescence was measured (ex: 530 nm, em: 590 nm) at 6 h post-addition of the solution to the cells. Results are from one of three independent experiments, data points in the graph are the mean of 6 technical replicates, error bars are standard deviation. **(C)** Table of N/P ratios at different combinations of pDNA and PEI quantities.

### *Evaluation of PEI toxicity on* A. castellanii *cells*

One factor often considered in transfection studies involving mammalian cells is how the transfection reagent affects the cells, as high concentrations can be toxic. Therefore, a balance between a high transfection rate and cytotoxicity should be struck. Here, a resazurin assay was performed to estimate the effect of PEI concentration on the amoebae. The use of this assay to evaluate *A. castellanii* viability has been described ^20^, but the assay was further optimised here to ascertain its sensitivity for the conditions used (**Figure S2**). Cellular toxicity varied between the PEIs, with the most prominent contrast between the linear and the branched PEIs (**Figure 1B**). PEI-40 had the least toxic effect on the *A. castellanii* cells, with only a 20% decrease in cell viability at the highest quantity tested (2.5 μg), followed by PEI-25 and PEI-Br with a 40% and 80% decrease, respectively. Altogether, the results suggest a threshold at a maximum of ∼1.5 μg of PEI, whereafter the most substantial decrease in viability is seen.

### Optimisation of transfection conditions

Transfection efficiency is highly dependent on the effective formation of DNA-PEI polyplexes, i.e., all the DNA should be condensed by the PEI so it can be delivered to the cell. For this to happen, there should be enough available nitrogen (N) in the PEI to cationise the phosphate (P) in the DNA, this is referred to as the N/P ratio, and it is used to find the optimal conditions for transfection. Since no previous work has described PEI-based transfection of *Acanthamoebae*, a systemic approach was employed to evaluate the performance of each PEI used in this study. A combinatorial matrix using a range of pDNA and PEI amounts resulted in an encompassing assay of 100 different combinations, with N/P ratios ranging from ∼ 0.7 to 70 (**Figure 1C**).

### Transfection efficiency is dependent on minimum quantities of pDNA and PEI

The results of the combinatorial transfection screen for the two constitutive fluorescence plasmids pGAPDH-EGFP and pGAPDH-mScarlet (**Figure S3**) illustrate a significant impact of PEI choice for transfection of *A. castellanii*. Overall, there is a minimum threshold for efficient transfection, contingent on the concentration of PEI. The results for PEI-Br concur with the toxicity data, as there were few transfectants at PEI quantities above 1.5 μg, with most fluorescent cells observed at N/P ratios ranging from ∼ 2.9 - 4.5 (**Figure 1C and Figure 2**).

**Figure 2.**
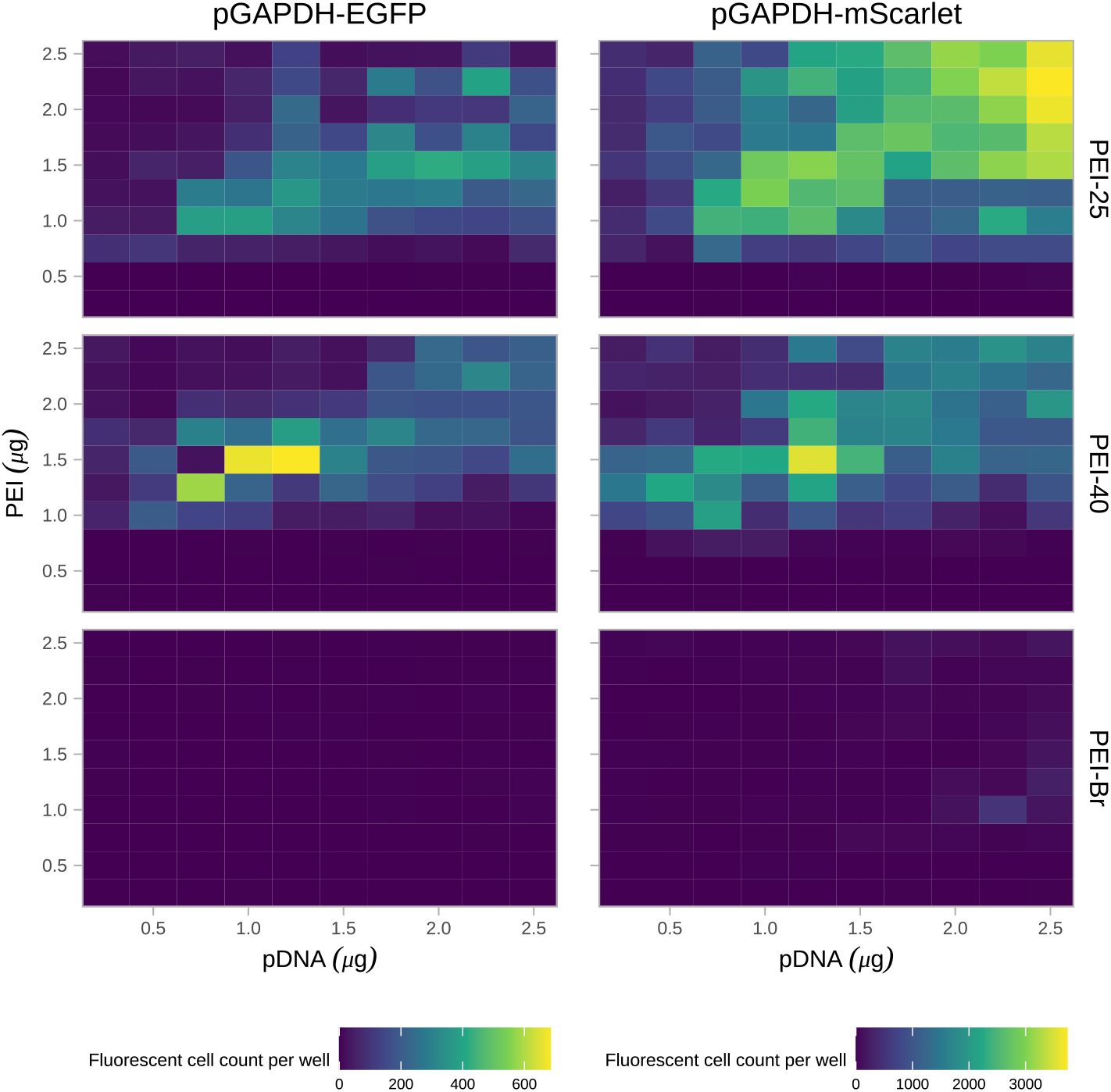
Effect of DNA and PEI quantities on transfection efficiency. The left and right panels show the results for cells transfected pGAPDH-EGFP and pGAPDH-mScarlet, respectively. The top, middle and bottom panels correspond to transfections with PEI-25, PEI-40 and PEI-Br, respectively. The x- and y-axis represent the amount of DNA and PEI, respectively. The colour of the tiles represents the sum of fluorescent cells detected per well. Note that scales are different for the left and right panels.

The linear PEIs (PEI-25 and PEI-40) had the highest transfection efficiencies, but the results were not directly correlated with higher N/P ratios. Although the results varied considerably between the two plasmids, the overall distribution trends were similar. For PEI-25 and PEI-40, the minimum threshold of PEI per reaction appeared to be above 0.5 μg and 0.75 μg PEI per reaction, respectively. Over this threshold, there were transfectants even with the lowest amounts of pDNA. The highest transfection efficiencies were observed at low to modest N/P ratios, i.e., ∼5 – 10 (**Figure 2**). Interestingly, the highest fluorescent cell count with PEI-40 for both plasmids was at the same N/P ratio of 8.6, and with PEI-25, there was a high number of fluorescent cells even at higher amounts of PEI.

### *Fluorescence intensity varies between* A. castellanii *transfectants*

Through microscopy, we observed that the level of fluorescence varied considerably between transfectants (**Figure 3** and **S4**). A notable variation in individual cell fluorescence appeared throughout all transfected cells (Figures **3A** and **3B**), but this was most pronounced on pGAPDH-mScarlet transfectants and particularly at high amounts of pDNA and PEI (**Figures 3C** and **3D**).

**Figure 3.**
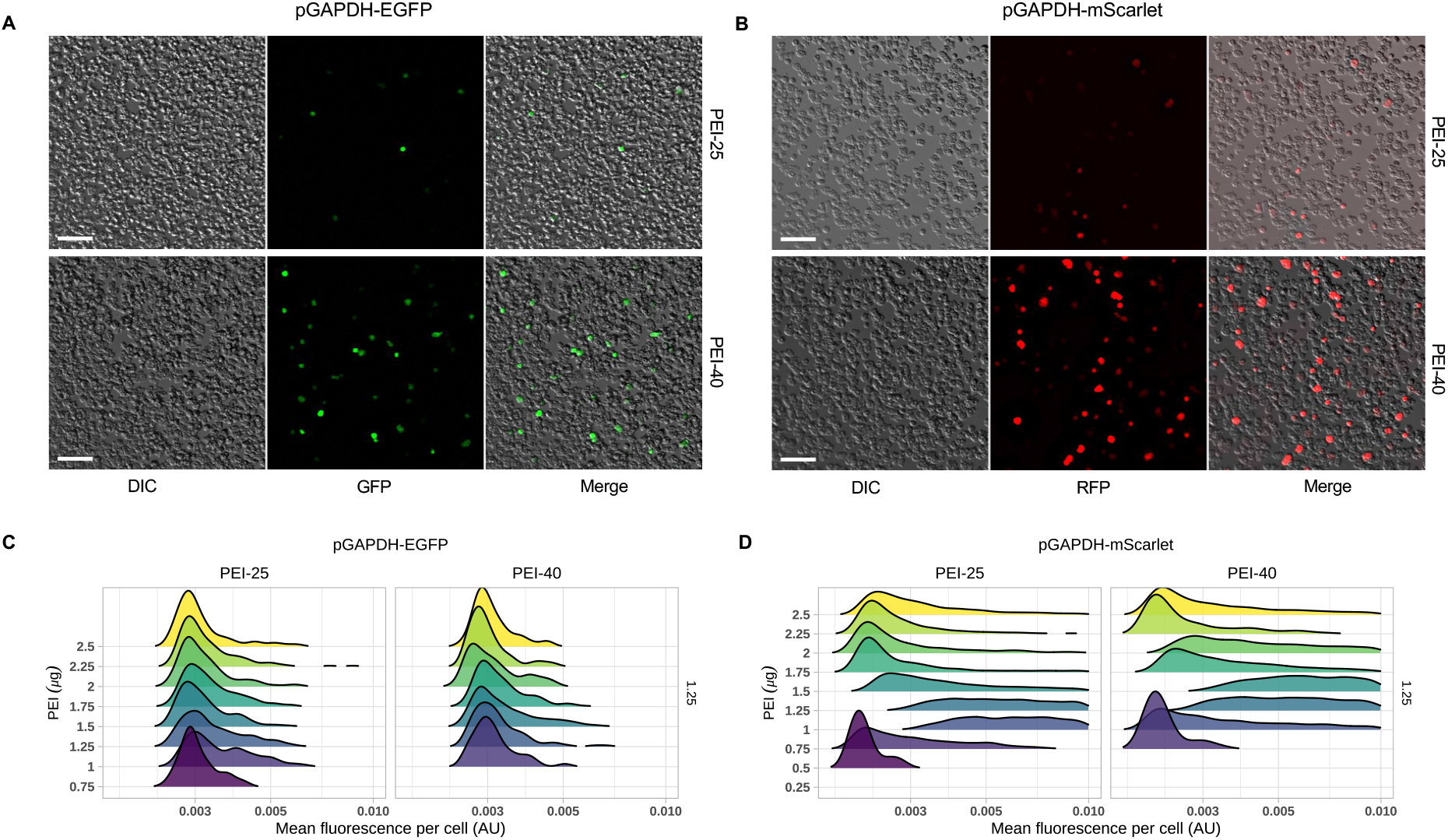
Individual fluorescence variation of *A. castellanii* transfectants. Representative micrographs of *A. castellanii* transfected with pGAPDH-EGFP (**A**) or -mScarlet (**B**) using PEI-25 or PEI-40. Scale bars, 100 µm. Mean fluorescence distribution per cell upon transfection with PEI-25 or PEI-40 and 1.25 μg of pGAPDG-EGFP (**C**) or pGAPDH-mScarlet (**D**). Each ridge corresponds to rising concentrations of PEI (bottom to top) within each panel, marked with different colours, showing the distribution of the cell fluorescence per condition (x-axis, logarithmic scale).

To discern whether the fluorescence variability was due to the number of plasmids per cell or to differential gene expression the amoebae were then transfected with both plasmids simultaneously. From the transfection screen results, it was clear that pGAPDH-mScarlet yields more transfectants than pGAPDH-EGFP, so it was imperative to optimise the ratio of pGAPDH-EGFP:pGAPDH-mScarlet plasmids per transfection reaction. For this analysis, the *A. castellanii* cells were co-transfected with PEI-40 at the optimal pDNA:PEI ratios (1.25:1.5 μg, as experimentally observed; **Figure 2**); and using a range of pGAPDH-EGFP:pGAPDH-mScarlet plasmid ratios from 10:0 to 0:10 (**Figure S5**).

As expected, pGAPDH-mScarlet yielded more transfectants also in this setup. Most importantly, at a 9:1 pGAPDH-EGFP:pGAPDH-mScarlet plasmid ratio, we observed a substantial fraction of EGFP+ and mScarlet+ co-transfectants (**Figure 4** and **S5**). This experiment illustrates that *A. castellanii* can be readily co-transfected with more than one plasmid per cell with the current protocol and that titration of protein expression levels can also be accomplished by altering pDNA amounts.

**Figure 4.**
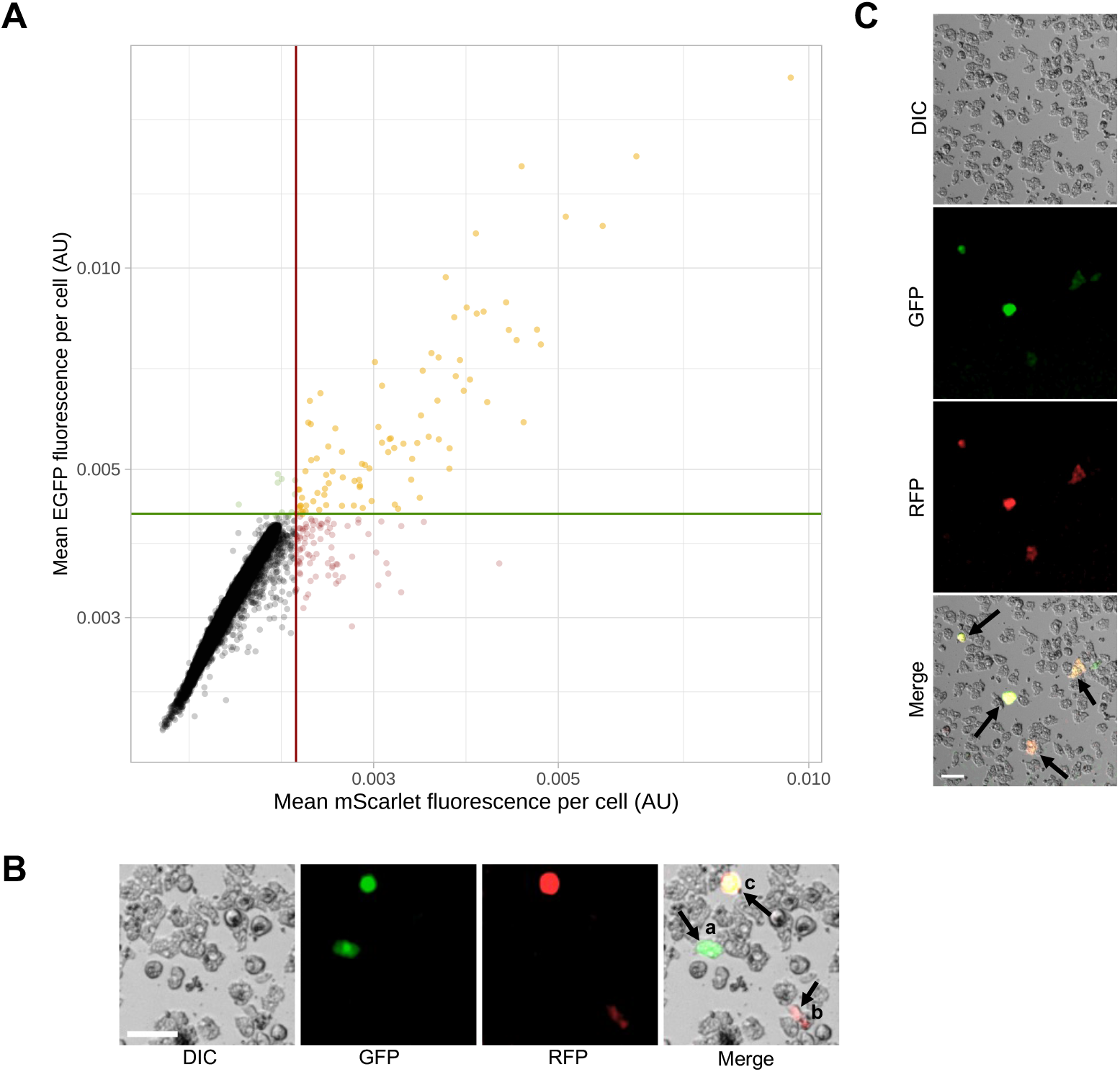
Distribution of transfectants according to their fluorescence. Co-transfection of *A. castellanii* at a 9:1 pGAPDH-EGFP:pGAPDH-mScarlet plasmid ratio. (**A**) Black dots are non-transfected cells, red and green are pGAPDH-EGFP and -mScarlet single-plasmid transfectants, respectively, and yellow are transfectants harbouring both plasmids. The x- (red/mScarlet) and y-intercept (green/EGFP) lines represent the minimum fluorescence threshold for a cell to be considered a positive transfectant. (**B**) Representative micrographs of transfected *A. castellanii*, arrows point to cells with plasmids expressing (a) EGFP, (b) mScarlet, and (c) both. (**C**) Representative micrographs of *A. castellanii* co-transfectants with varying EGFP and mScarlet-fluorescence intensities. Scale bars, 50 µm.

### The use of PEI greatly improves transfection efficiency over currently used reagents

The efficiency of PEI as a transfection reagent for *A. castellanii* was next compared to that of the currently used commercial reagents, namely SuperFect^®^ and ViaFect™. A common transfection protocol for each reagent was derived from published studies on *A. castellanii* ^11,12^. The two resulting protocols differed in that transfection was done with 1 µg of pDNA for SuperFect^®^, and 0.2 µg for Viafect™, and incubation times of the complexes with cells were 3 h for SuperFect^®^ and 16 h for Viafect™. The two protocols were then adapted for PEI-25 and -40 (i.e., using the same quantity of pDNA and incubation period) in combination with the optimal quantity of each PEI for these pDNA concentrations (informed by results in **Figure 2**). The outputs of each protocol using the original reagent were then compared to PEI-25 and -40 and quantified using CellProfiler, as before (**Figure 5**).

**Figure 5.**
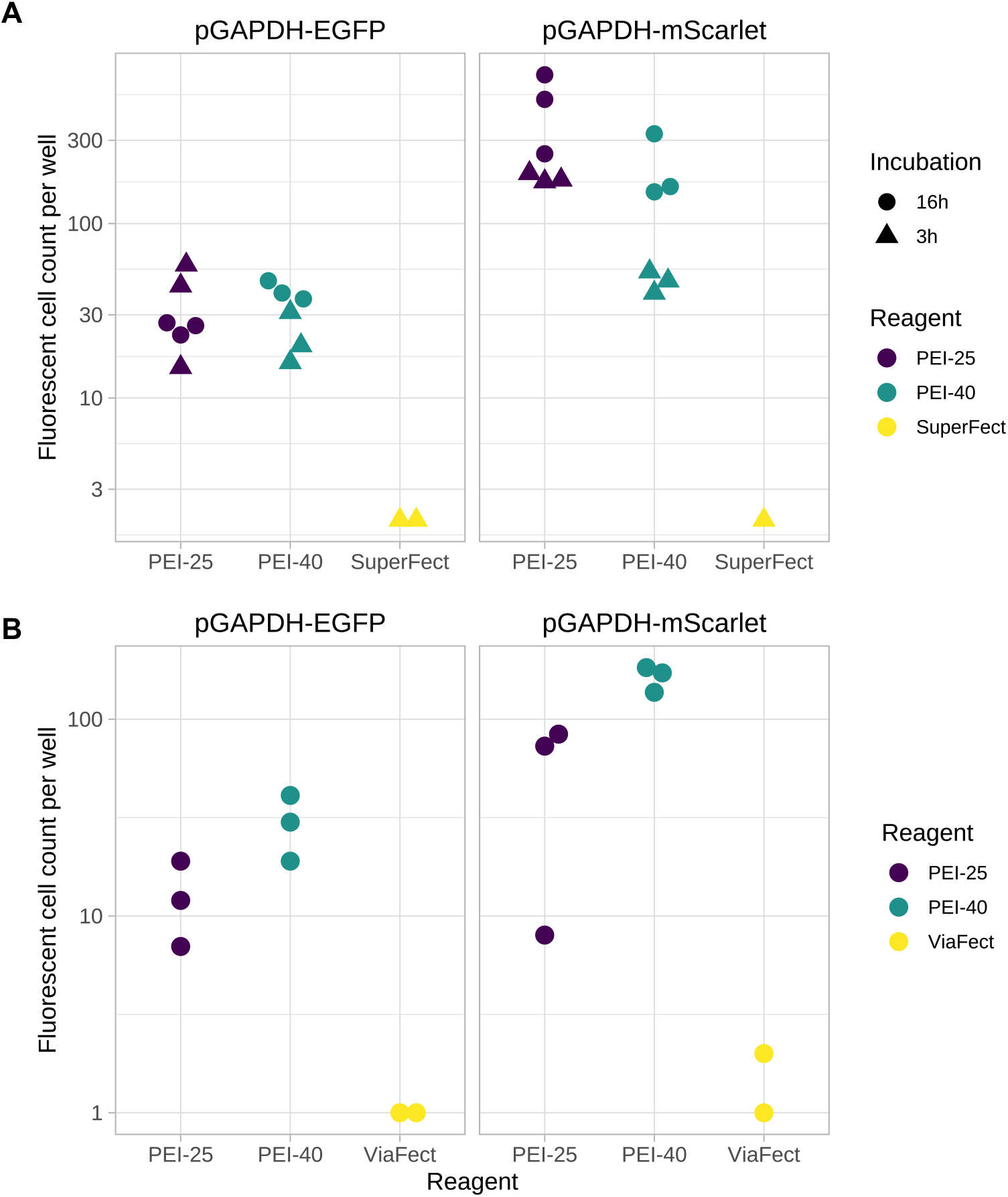
Comparison of the transfection efficiency of PEI-25 and 40 *vs* SuperFect^*®*^ or Viafect™ during 3 h or 16 h incubation. (**A**) Fluorescent cell count per well after transfection of *A. castellanii* with pGAPDH-EGFP or -mScarlet using PEI-25, PEI-40 or SuperFect^*®*^. Per manufacturer instructions pDNA:SuperFect^*®*^ complexes were incubated with the cells for 3 h, while pDNA:PEI complexes were incubated with the cells for 3 h or 16 h. The symbol colours correspond to the different reagents and the y-axis to the total transfectant count. Each symbol is a technical replicate. Experiments were conducted in triplicates. Some experiments with SuperFect^*®*^ or Viafect™ did not yield any transfectant and are thus not displayed. The triangle and dot shapes represent the incubation period of the complexes with the cells (3 and 16 h, respectively). (**B**) shows the same comparison between PEIs and Viafect™.

In both comparisons, the PEIs were vastly more efficient at transfecting the amoebae than either commercial reagent (**Figure 5**). Compared to SuperFect^®^, both PEIs were at least 5x to up to ∼100x more efficient when using PEI-25 and pGAPDH-mScarlet (**Figure 5A**). In the PEIs *vs* Viafect™ comparisons, although the relative data was similar to SuperFect^®^, the absolute number of transfectants was lower on average, which is congruent with this study’s previous results from transfections with low pDNA amounts (**Figure 2**). Also here, both PEIs were more efficient, at 8 to over 100-fold the transfection efficiency of Viafect™ (**Figure 5B**).

As the transfections using PEIs had not been tested with shorter incubation periods, replica transfections were performed in which the *A. castellanii* were incubated with complexes for 3 h (**Figure 5A, triangles**). These results indicate that pDNA:PEI complexes can be transfected with broadly similar efficiency even using significantly shorter incubation periods. In summary, PEI-25 and -40 vastly outperform SuperFect^®^ and ViaFect™ even under (for PEI) sub-optimal conditions.

## Discussion

*A. castellanii* is a free-living protist, ubiquitous and abundant in freshwater and soil habitats. It is one of the most often isolated amoebal species worldwide ^21,22^. Its status as an emerging human pathogen is due not only to the infections it causes as an opportunistic human pathogen but also because of its role as a reservoir for bacteria, viruses, and fungi ^21,22,23^. Furthermore, due to the cellular and functional similarities between *A. castellanii* and mammalian phagocytic cells, the former can be considered training grounds for these pathogens to cause human infection ^21,22,24^. These factors make *A. castellanii* an important and exciting model organism. Still, due to the shortage of molecular biology techniques (among others), it has not been as widely used as other amoebae ^21–24^.

Here, we demonstrated that the readily available and inexpensive cationic polymer PEI can be used to efficiently transfect *A. castellanii* with plasmid DNA. We systematically explored the use of three PEIs, PEI and DNA quantities, on transfection efficiency and their effects on cell viability. Our combinatorial approach allowed us to test a broad range of PEI and pDNA amounts and ratios in a high-throughput manner. The N/P ratio range tested was sufficient to highlight how there are none or very few transfectants at the extremes of the N/P ratio distribution. It also highlights how the actual quantity of the reagents in the solution, and not just the N/P ratio, plays an important role in efficient transfection.

These observations provide a comprehensive guide to choosing optimal transfection parameters for *A. castellanii*.

We also demonstrate that co-transfection of more than one plasmid per cell is possible and prevalent. Furthermore, the quantity of PEI used affected not only the absolute transfection efficiency, but also the distribution of the different transfected plasmids during co-transfection, making it possible to modulate the level of co-transfection. This could be used to normalise or titrate the number of each plasmid transfected per cell, should it be desirable to have a homogenous or a graded protein expression within the population. However, the differing transfection efficiency between the two fluorescence plasmids remains unclear. Two hypotheses may explain this phenomenon: (i) mScarlet is a brighter fluorescent molecule, compared to EGFP (brightness of 70 vs 33.54, respectively, according to www.fpbase.org), and so the plasmids may be similarly transfected, but mScarlet fluorescence may be more easily detected; alternatively, (ii) while EGFP was codon-optimised for expression in humans, the mScarlet sequence was codon optimised for *A. castellanii*, which should improve the expression of the latter. This effect has been described before, e.g. Bateman observed that: “*(…) high levels of monomeric red fluorescent protein (mRFP) expression could be serendipitously obtained*.” when using pTPBF-mRFP *versus* pTPBF-EGFP ^25^. Another report mentions using a 1.7 ratio of mVenus-over tdTomato-expressing plasmid when co-transfecting to account for differences in brightness and quantum yield of the fluorescent proteins ^26,27^. Temperature, an element to consider when choosing fluorescent proteins ^27^, might also provide some insight into this phenomenon; since EGFP has been optimised for expression in human cells, it stands to reason that this protein would also be best expressed at 37°C and not at 30°C used here. Ultimately, no generalisable conclusions regarding if one plasmid is always *better* than another should be drawn from the present work. Nevertheless, we demonstrate that PEI transfection provides a vastly improved protocol over current commercial standards, both for an EGFP and a mScarlet fluorescence plasmid.

The cytotoxicity results show that even high concentrations of PEI-25 and PEI-40 are non-toxic. Yet, it is important to be wary that high concentrations of PEI could affect the cells in other ways not assessed here. Therefore, other cell characteristics, such as morphology, should be considered in future experiments. Also not investigated here, but of interest, is whether PEI is also effective for transfection of siRNA in *A. castellanii*. Many studies demonstrate PEI-mediated gene knockdown with siRNA in other cell types. In studies where *A. castellanii* is transfected with siRNA the reagents used are the same as those used for transfection with plasmid DNA ^12^, so it can be surmised that PEI would also work to achieve this in *A. castellanii*.

In summary, we here provide evidence for a high-efficiency transfection protocol in *A. castellanii* using the cationic polymer PEI and comprehensively optimise the relevant transfection conditions.

## Materials and Methods

### Acanthamoeba *culture*

*Acanthamoeba castellanii* (ATCC 30010) was cultured in PYG medium with additives (ATCC medium 712) at 30°C, and cultures were split three times per week.

### Plasmid DNA

The pGAPDH-EGFP plasmid used in this study was first described by Bateman ^25^, and it was kindly sent to us by Prof. Yeongchul Hong (Kyungpook National University, Korea) (**Figure S3**). To make pGAPDH-mScarlet (**Figure S3**), the EGFP fluorescence reporter gene was removed by enzymatic digestion of the plasmid with *Nde*I and *Xba*I restriction enzymes. The plasmid backbone was gel purified and used for ligation with mScarlet cut with the same restriction enzymes as above. The sequence of the mScarlet gene ^28^ was codon optimised according to the codon profile of *A. castellanii*, using a codon optimisation tool (https://dnahive.fda.gov/dna.cgi?cmd=codon_usage&id=537&mode=cocoputs) ^29^. The restriction sites – *Nde*I and *Xba*I – were added at the 5’ and 3’ ends, and Integrated DNA Technologies® synthesised the sequence.

### Transfection reagents

The 25kDa PEI 25™ (PEI-25) and 40 kDa PEI Max^®^ (PEI-40) polymers were acquired from Polysciences and were prepared according to manufacturer instructions to get a stock solution of 1 mg/ml. Branched PEI (PEI-Br) was purchased from Sigma-Aldrich (408727) and diluted in ultrapure water to 1 mg/ml concentration.

### Plasmid DNA-PEI affinity in different media

To test the effect of different media and buffers on the ability of the PEI to form complexes with the plasmid DNA at different N/P ratios, increasing amounts of PEI (0.25 – 2.5 μg) and 0.25 μg of pDNA were each diluted in 10 μl of the indicated complexation buffers, mixed by pipetting and incubated at RT for 20 minutes. After incubation, 30 μl of the respective buffer was added to the pDNA-PEI complex mix. An aliquot of this was used to load a 0.7% agarose gel (with GelRed), and gel electrophoresis was carried out for 1h at 100V. The different media used were Peptone Yeast Glucose (PYG) (ATCC 712), Ac medium (PYG medium without protease peptone, yeast extract, and glucose), and LoFlo (ForMedium); and HEPES buffered solution (HBS), NaCl (150 mM) solution, and ultrapure water (ddH_2_O).

### Viability Assay standardisation

To assess the toxicity of the transfection reagents, viability assays were performed using a resazurin assay ^20^. This assay reduces the non-toxic and non-fluorescent dye resazurin to fluorescent resorufin via mitochondrial reductase. Changes in fluorescence indicate mitochondrial activity, a proxy for cell viability. Resazurin sodium salt (Sigma) was diluted in PBS to 2 mM, filter-sterilised, and kept at 4°C, protected from light.

To determine the optimal cell number and incubation period with the resazurin salt solution for the assay, 2.5×10^3^-8.0×10^4^ *A. castellanii* cells were seeded per well, in triplicates, in a black 96-well plate, in a final volume of 100 μl of PYG. The plate was incubated at 30°C for 16 h. Then, 10 μl of the resazurin solution was added to each well, the plates were further incubated, and fluorescence measurements were taken at 2, 4 and 6 h. The resorufin fluorescence was measured at 530 nm (excitation) and 590 nm (emission) at 5 locations in the well using a Tecan Spark fluorometer^®^. Measuring the fluorescence produced by the amoebae with a 2 mM resazurin solution and 6 h of incubation proved sensitive enough to discriminate cell viability even at low cell concentrations.

### PEI toxicity assays

*A. castellanii* tolerance to each PEI reagent was assessed using the resazurin assay. Briefly, 2×10^4^ cells were seeded per well in a black 96-well plate and incubated overnight at 30°C. The following day the medium was replaced by 50 μl of PYG containing increasing concentrations of PEI (0.25-2.5 μg), and the plates were incubated for 24 h. The medium was removed, and 100 μl PYG was added before initiating the resazurin assay (described above).

### Nitrogen/Phosphate ratio theoretical approximation

The ratio of the amount of nitrogen in PEI per phosphate in DNA in a reaction mixture (N/P ratio) was estimated, as it plays an essential role in polyplex formation and transfection efficiency. The theoretical amine (nitrogen) groups in PEI are derived from the polymer’s chemical formula for a single repeat unit (monomer). Using the 25kDa lPEI (-NHCH_2_CH_2_-) as an example, a monomer’s molecular weight (MW) would be 43 g/mol. Therefore, 1 μg would contain 23 nmol of nitrogen. The number of phosphate groups in the DNA is calculated using the equations (1) and (2)^13,14,18^:

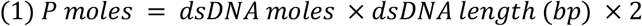

Where

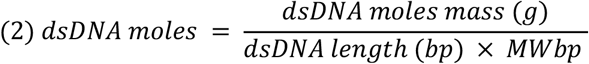

Hence, 1 μg pGAPDH-EGFP (5831 bp) would have 3.2 nmol of phosphate. The N/P ratio of this reaction is then calculated with the following equation (3):

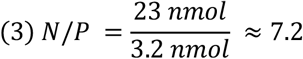

### Acanthamoeba *transfection*

A day before transfection, 1×10^4^ cells were seeded per well in a 96 black well plate, in 100 μl of PYG with 1% Penicillin-Streptomycin (PS). On the day of transfection, pDNA and PEIs were each first diluted in 10 μl of PYG, then mixed by pipetting, and incubated at room temperature for 20 mins. During this incubation, the amoebae plates were prepared by aspirating the PYG, and washing once. Lastly, 30 μl of PYG was added to the pDNA-PEI mix, the total volume (50 μl) was to each well, drop-wise, and the plates were then incubated at 30°C for 16 h.

### Transfection with SuperFect^®^ and Viafect™

To compare PEI-25 and PEI-40 and the commercial reagents – Viafect™ and SuperFect^®^, *A. castellanii* cells were prepared and seeded as described above. The ratios of pDNA to transfection reagent were based on published methods ^11,12^ and the manufacturer’s protocols. For transfections with SuperFect, 1 µg pDNA was mixed in 30 µl of PYG, before adding 5 µl of SuperFect^®^. This reaction was incubated at room temperature for 20 mins, then 150 µl of PYG was mixed in and added to the cells. The cells were then incubated for 3 h at 30°C, after which the DNA-SuperFect^®^ solution was carefully aspirated, the cells were washed once, and fresh PYG with PS was added, followed by overnight incubation. For the paired comparison assays, 1 µg pDNA was transfected with 1.5 µg of PEI-25/PEI-40, as described, cells were incubated with the DNA: PEI complexes, for 3 h or 16 h, before replacing the medium with PYG with PS. Viafect™ transfections were performed similarly: 0.2 µg pDNA and 1 µl reagent were mixed in 50 µl PYG; this solution was incubated with the cells for 3 h, before adding 100 µl of PYG and further incubating for a total of 16 h. To compare the PEIs with Viafect™, 0.2 µg pDNA was mixed with 1 µg PEI-25/PEI-40 and incubated with the cells for 16 h.

### Detection of transgene expression

To assess the efficiency of transfection, a microscopy-based quantification approach was taken. Using a Nikon T*i*-2 Eclipse microscope equipped with a Spectra-X light engine (Lumencor), 4 pictures were taken at different locations per well, with a 10x /0.45 NA Plan Apochomat objective and a Photometrics Prime 95B camera (1.12 µm pixel size), covering a total area of approximately 38% of the well. Micrographs were taken using DIC contrast and in the GFP (excitation: 475/34, emission filter: Chroma ET510/20) or RFP (excitation: 575/35, emission Chroma 89403) channels. The images were segmented using the Cellpose 2.0 segmentation algorithm ^30^, processed and analysed in CellProfiler v4.2.1^31^. Data were further analysed with R ^32^, and figures were produced with the R package ggplot2 ^33^ and GraphPad Prism (version 9.4.0 for Mac).

## Acknowledgements

We thank Erik Wistrand-Yuen for his help in plasmid construction and Claudia Bergin and Anna-Maria Divne, from the SciLifeLab Microbial Single Cell Genomics Facility at Uppsala University, for their assistance with the initial fluorescence microscopy.

## Author contributions

A.B.M.: Conceptualisation, Methodology, Investigation, Formal Analysis, Visualisation, Writing - Original draft. V.E.: Visualisation, Software. J.E.: Methodology, Software. M.E.S.: Methodology, Resources. L.G.: Formal Analysis, Visualisation, Resources. All authors reviewed and edited the manuscript.

## Declaration of interests

The authors declare no competing interests.

## Supplementary Figures

**Figure S1.**
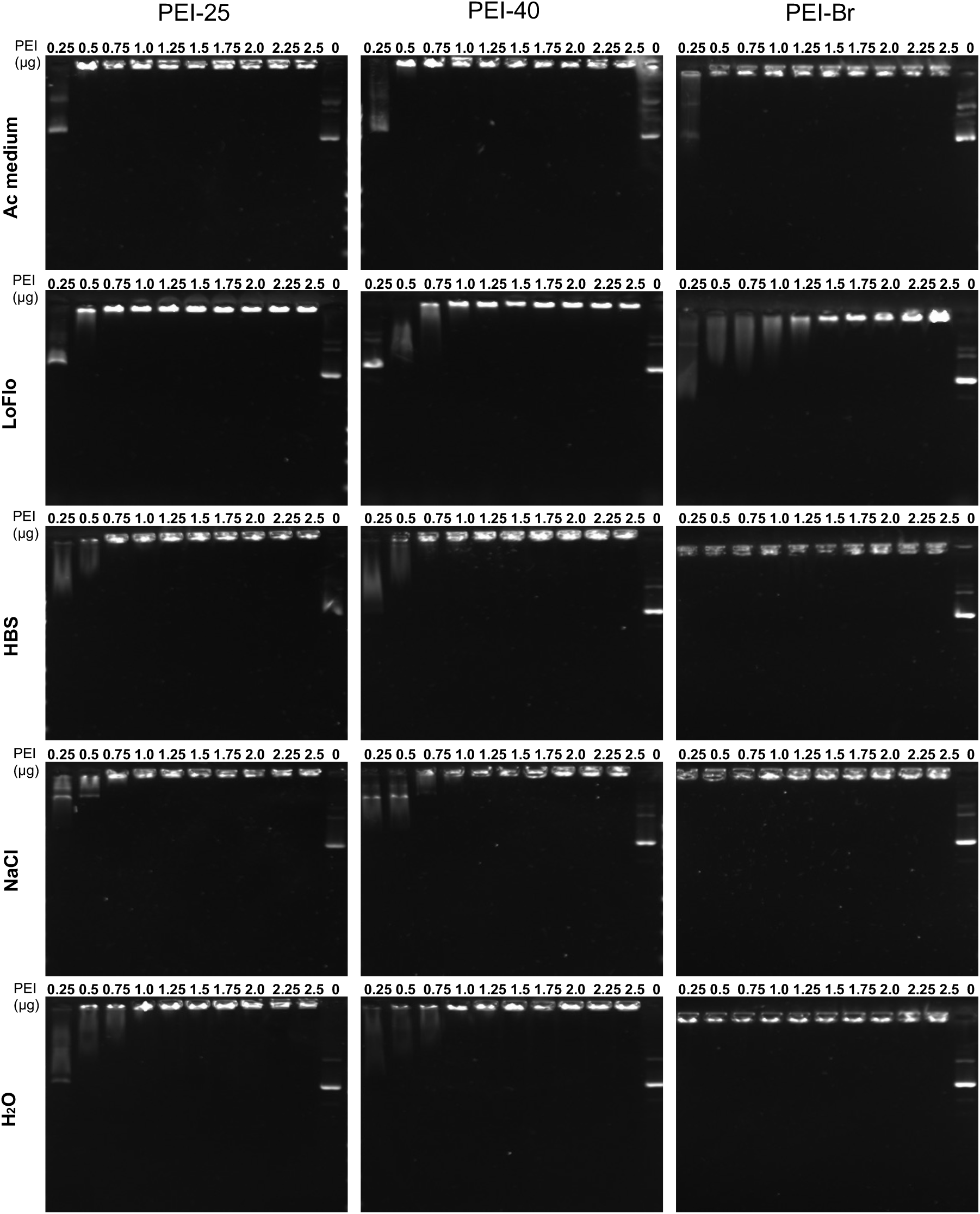
Condensation of pDNA with PEIs in different media. Increasing concentrations of PEI were used to form complexes with 0.25 μg of pDNA. The migration of the DNA-PEI polyplexes was compared to that of non-complexed/naked pDNA (C, untreated control).

**Figure S2.**
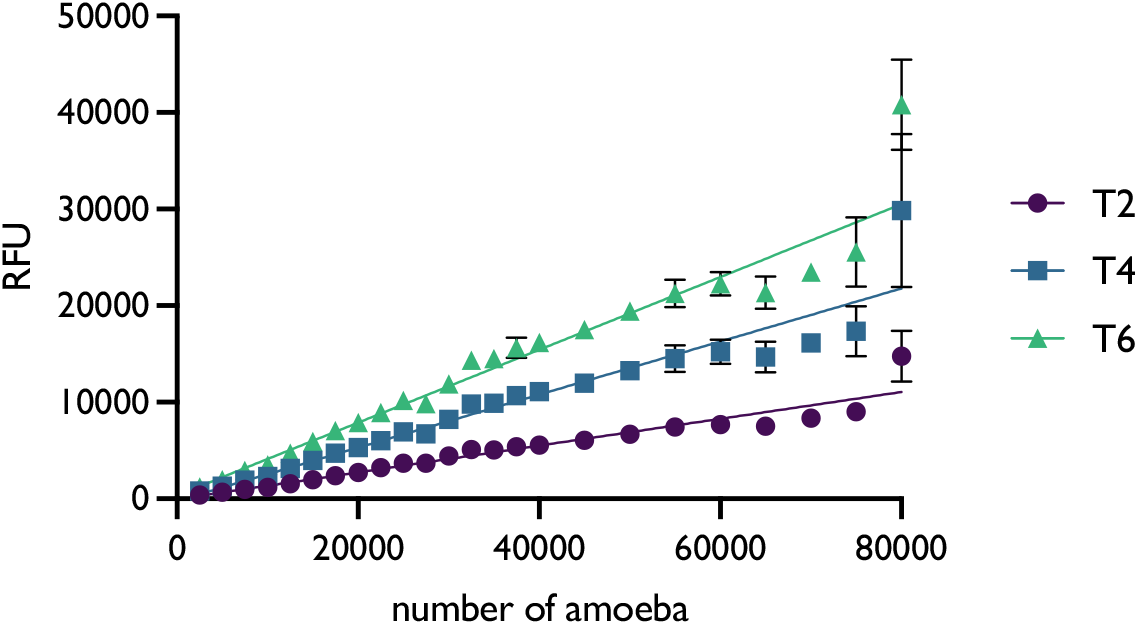
Resazurin viability assay optimisation. Fluorescence signal produced by *A. castellanii* using 2 mM resazurin sodium salt solution at different incubation periods (2, 4 and 6 hours). Results are the mean of 6 technical replicates and 3 independent experiments; error bars are standard deviation.

**Figure S3.**
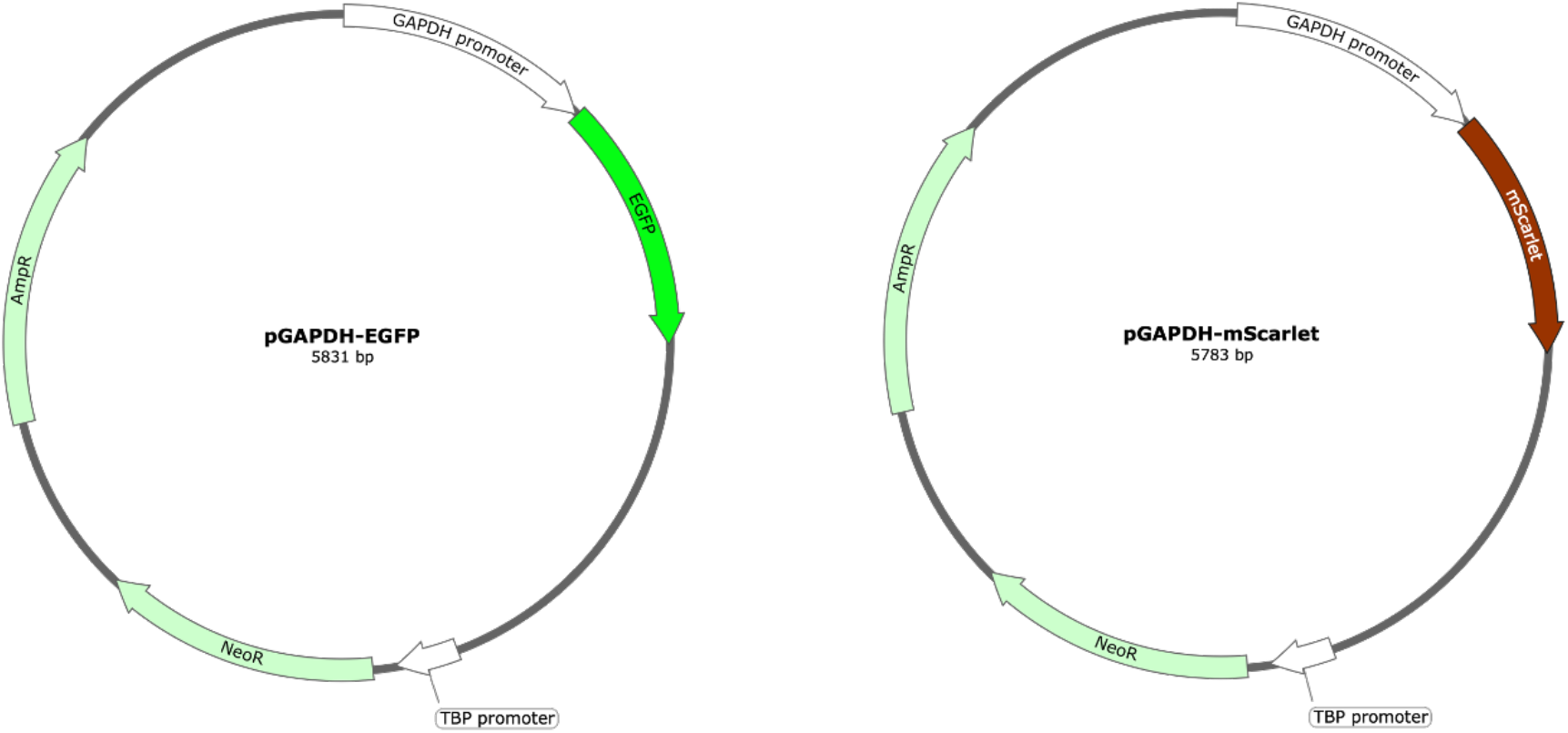
Plasmids used in this study. Plasmid pGAPDH-EGFP was first described in Bateman (2010) ^25^, and plasmid pGAPDH-mScarlet was constructed as part of this study. Both plasmids contain the *Acanthamoeba* GAPDH gene promoter upstream of the fluorescence marker (EGFP or mScarlet) for constitutive expression in *A. castellanii*; an ampicillin resistance gene (AmpR) for growth and selection in *E. coli*; a neomycin resistance gene (NeoR) downstream of the *Acanthamoeba* TBP gene promoter for selection in *A. castellanii*.

**Figure S4.**
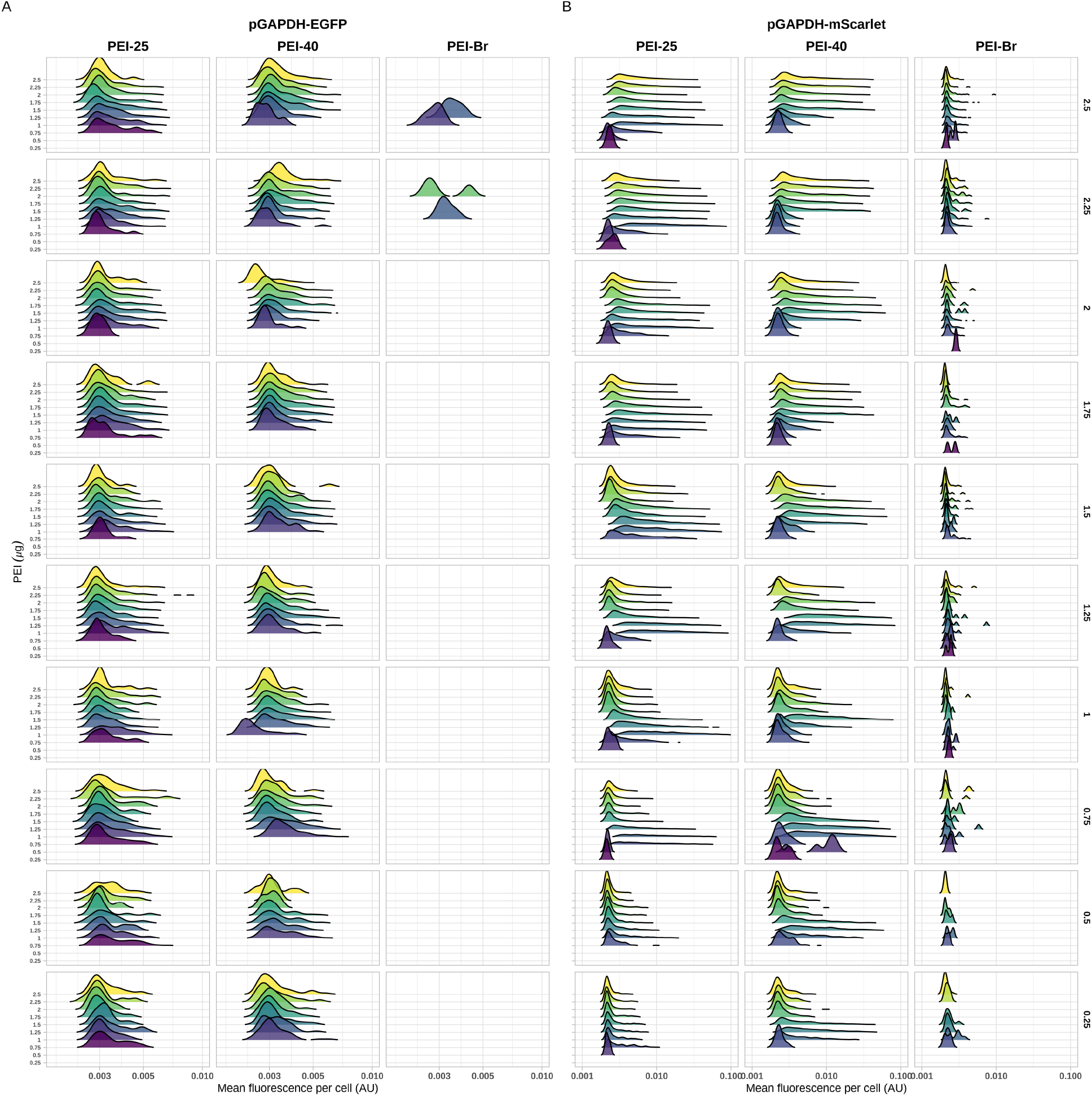
Fluorescence distribution per cell transfected with different pDNA and PEI quantities. Transfection of *A. castellanii* with pGAPDH-EGFP (**A**) or pGAPDH-mScarlet (**B**) plasmids using PEI-25, -40 and -Br. The rows of panels correspond to, from top to bottom, decreasing concentrations of pDNA. In each panel, the ridges illustrate the fluorescence distribution per cell with increasing quantities of PEI.

**Figure S5.**
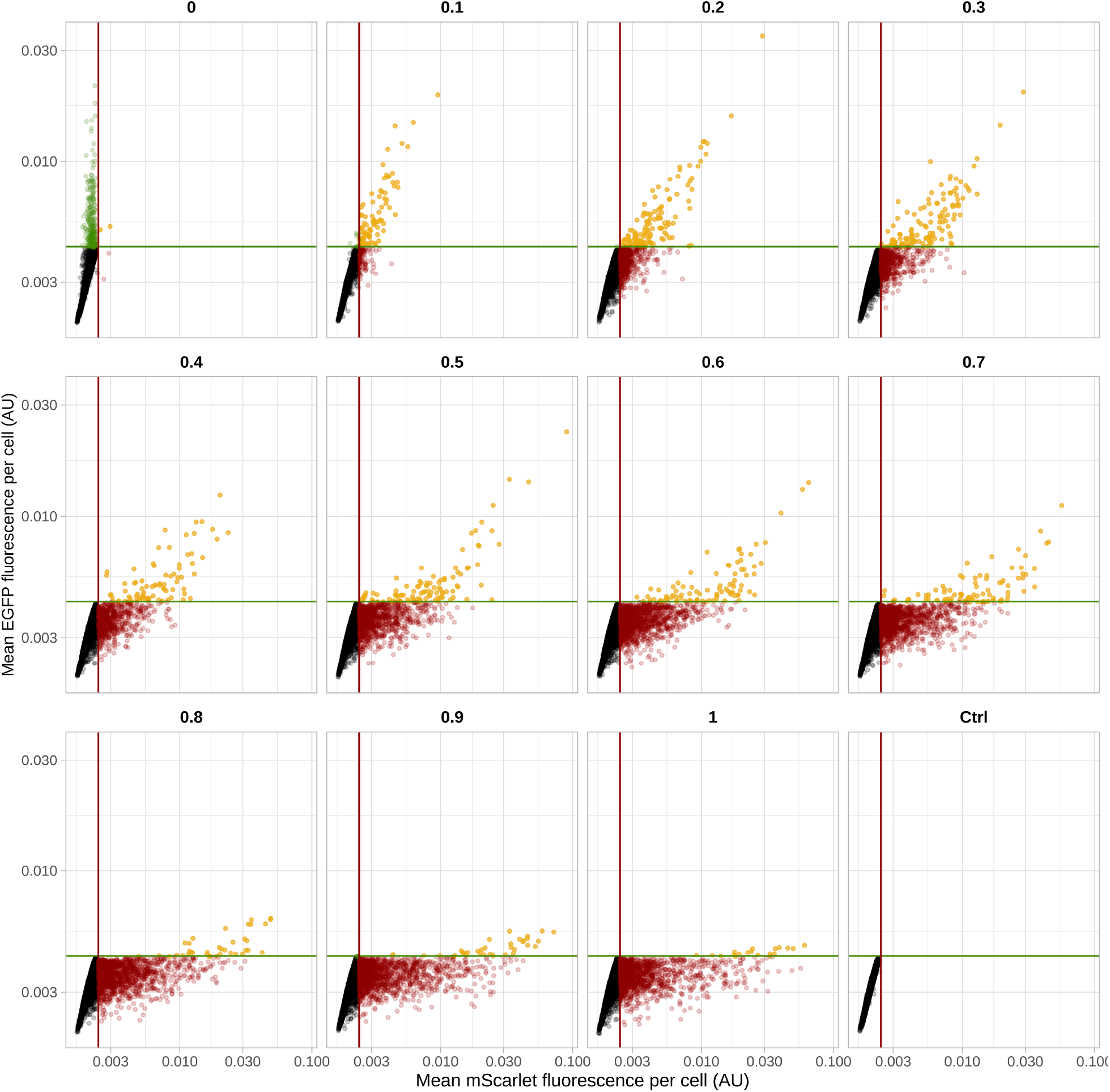
Distribution of *A. castellanii* co-transfectants according to their fluorescence. Transfection of *A. castellanii* with different ratios of pGAPDH-EGFP and pGAPDH-mScarlet plasmids (indicated above each plot). In each plot: black dots are non-transfected cells; red and green dots are pGAPDH-EGFP and -mScarlet-single plasmid transfectants, respectively; and yellow dots are co-transfectants with both pGAPDH-EGFP and -mScarlet plasmids. The X- (red/mScarlet) and Y-intercept (green/EGFP) lines represent the threshold fluorescence for a cell to be considered a positive transformant. The ‘Ctrl’ plot is non-transfected cell control.

## Notes

### Competing Interest Statement

The authors have declared no competing interest.

### Summary of Updates

In this revision, we condensed the text in the results part. We combined some figures together: Fig. 1, 2 and Table 1 became Fig. 1; Fig. 3 became Fig. 2; Fig. 4 was partly moved to the supplementary, and images from cultures were added to the main manuscript (Fig. 3); Fig. 5 became Fig. 4, with added images from fluorescent cells; Fig. 6 became Fig. 5.

## References

1. Greub, G. & Raoult, D. Microorganisms Resistant to Free-Living Amoebae. Clinical Microbiology Reviews 17, 413–433 (2004).

2. Peng, Z., Omaruddin, R. & Bateman, E. Stable transfection of Acanthamoeba castellanii. Biochimica et Biophysica Acta - Molecular Cell Research 1743, 93–100 (2005).

3. Swart, A. L., Harrison, C. F., Eichinger, L., Steinert, M. & Hilbi, H. Acanthamoeba and Dictyostelium as Cellular Models for Legionella Infection. Frontiers in Cellular and Infection Microbiology 8, (2018).

4. Siddiqui, R. & Khan, N. A. Biology and pathogenesis of Acanthamoeba. Parasites & Vectors 5, 6 (2012).

5. Clarke, M. et al. Genome of Acanthamoeba castellanii highlights extensive lateral gene transfer and early evolution of tyrosine kinase signaling. Genome Biology 14, R11 (2013).

6. Matthey-Doret, C. et al. Chromosome-scale assemblies of Acanthamoeba castellanii genomes provide insights into Legionella pneumophila infection–related chromatin reorganization. Genome Res. 32, 1698–1710 (2022).

7. Moon, E.-K. et al. Construction of EST Database for Comparative Gene Studies of Acanthamoeba. Korean J Parasitol 47, 103–107 (2009).

8. Samba-Louaka, A. Encystment of Free-Living Amoebae, So Many Blind Spots to Cover. Parasitologia 3, 53–58 (2023).

9. Leitsch, D., Mbouaka, A. L., Köhsler, M., Müller, N. & Walochnik, J. An unusual thioredoxin system in the facultative parasite Acanthamoeba castellanii. Cell. Mol. Life Sci. 78, 3673–3689 (2021).

10. Kong, H.-H. & Pollard, T. D. Intracellular localization and dynamics of myosin-II and myosin-IC in live Acanthamoeba by transient transfection of EGFP fusion proteins. Journal of cell science 115, 4993–5002 (2002).

11. Lee, Y.-R. et al. Essential Role for an M17 Leucine Aminopeptidase in Encystation of Acanthamoeba castellanii. PLOS ONE 10, e0129884 (2015).

12. Rolland, S. et al. Encystment Induces Down-Regulation of an Acetyltransferase-Like Gene in Acanthamoeba castellanii. Pathogens 9, 321 (2020).

13. Boussif, O. et al. A versatile vector for gene and oligonucleotide transfer into cells in culture and in vivo: polyethylenimine. PNAS 92, 7297–7301 (1995).

14. Bono, N., Ponti, F., Mantovani, D. & Candiani, G. Non-Viral in Vitro Gene Delivery: It is Now Time to Set the Bar! Pharmaceutics 12, 183 (2020).

15. Mady, M. M., Mohammed, W. A., El-Guendy, N. M. & Elsayed, A. A. Effect of polymer molecular weight on the DNA/PEI polyplexes properties. Romanian Journal of Biophysics 21, 15.

16. Costa, D., Valente, A. J. M., Queiroz, J. A. & Sousa, Â. Finding the ideal polyethylenimine-plasmid DNA system for co-delivery of payloads in cancer therapy. Colloids and Surfaces B: Biointerfaces 170, 627–636 (2018).

17. Delafosse, L., Xu, P. & Durocher, Y. Comparative study of polyethylenimines for transient gene expression in mammalian HEK293 and CHO cells. Journal of Biotechnology 227, 103–111 (2016).

18. González-Domínguez, I., Puente-Massaguer, E., Lavado-García, J., Cervera, L. & Gòdia, F. Micrometric DNA/PEI polyplexes correlate with higher transient gene expression yields in HEK 293 cells. New Biotechnology 68, 87–96 (2022).

19. Song, K.-J. et al. Heat shock protein 70 of Naegleria fowleri is important factor for proliferation and in vitro cytotoxicity. Parasitol Res 103, 313 (2008).

20. Heredero-Bermejo, I. et al. In vitro comparative assessment of different viability assays in Acanthamoeba castellanii and Acanthamoeba polyphaga trophozoites. Parasitol Res 112, 4087–4095 (2013).

21. Rayamajhee, B. et al. Acanthamoeba, an environmental phagocyte enhancing survival and transmission of human pathogens. Trends in Parasitology (2022) doi:10.1016/j.pt.2022.08.007.

22. Guimaraes, A. J., Gomes, K. X., Reis Cortines, J., Peralta, J. M. & Peralta, R. H. S. Acanthamoeba spp. as a universal host for pathogenic microorganisms: One bridge from environment to host virulence. Microbiological Research 193, 30–38 (2016).

23. Molecular parasitology: protozoan parasites and their molecules. (Springer, 2016).

24. Hilbi, H., Weber, S. S., Ragaz, C., Nyfeler, Y. & Urwyler, S. Environmental predators as models for bacterial pathogenesis. Environmental Microbiology 9, 563–575 (2007).

25. Bateman, E. Expression plasmids and production of EGFP in stably transfected Acanthamoeba. Protein Expression and Purification 70, 95–100 (2010).

26. Colin, Béatrice, Deprez, Benoit, & Couturier, Cyril. High-Throughput DNA Plasmid Transfection Using Acoustic Droplet Ejection Technology. SLAS DISCOVERY: Advancing the Science of Drug Discovery 24, 492–500 (2019).

27. Shaner, N. C., Steinbach, P. A. & Tsien, R. Y. A guide to choosing fluorescent proteins. Nat Methods 2, 905–909 (2005).

28. Bindels, D. S. et al. mScarlet: a bright monomeric red fluorescent protein for cellular imaging. Nature Methods 14, 53–56 (2017).

29. Athey, J. et al. A new and updated resource for codon usage tables. BMC Bioinformatics 18, 391 (2017).

30. Stringer, C. & Pachitariu, M. Cellpose 2.0: how to train your own model. http://biorxiv.org/lookup/doi/10.1101/2022.04.01.486764 (2022) xdoi:10.1101/2022.04.01.486764.

31. Stirling, D. R. et al. CellProfiler 4: improvements in speed, utility and usability. BMC Bioinformatics 22, 433 (2021).

32. Team, R. C. R: A language and environment for statistical computing. R Foundation for Statistical Computing, Vienna, Austria. http://www.R-project.org/ (2016).

33. Wickham, H. ggplot2: Elegant Graphics for Data Analysis. (Springer New York, NY, 2009).

